# Microscopy-based assay for semi-quantitative detection of SARS-CoV-2 specific antibodies in human sera

**DOI:** 10.1101/2020.06.15.152587

**Authors:** Constantin Pape, Roman Remme, Adrian Wolny, Sylvia Olberg, Steffen Wolf, Lorenzo Cerrone, Mirko Cortese, Severina Klaus, Bojana Lucic, Stephanie Ullrich, Maria Anders-Össwein, Stefanie Wolf, Berati Cerikan, Christopher J. Neufeldt, Markus Ganter, Paul Schnitzler, Uta Merle, Marina Lusic, Steeve Boulant, Megan Stanifer, Ralf Bartenschlager, Fred A. Hamprecht, Anna Kreshuk, Christian Tischer, Hans-Georg Kräusslich, Barbara Müller, Vibor Laketa

## Abstract

Emergence of the novel pathogenic coronavirus SARS-CoV-2 and its rapid pandemic spread presents numerous questions and challenges that demand immediate attention. Among these is the urgent need for a better understanding of humoral immune response against the virus as a basis for developing public health strategies to control viral spread. For this, sensitive, specific and quantitative serological assays are required. Here we describe the development of a semi-quantitative high-content microscopy-based assay for detection of three major classes (IgG, IgA and IgM) of SARS-CoV-2 specific antibodies in human samples. The possibility to detect antibodies against the entire viral proteome together with a robust semi-automated image analysis workflow resulted in specific, sensitive and unbiased assay which complements the portfolio of SARS-CoV-2 serological assays. The procedure described here has been used for clinical studies and provides a general framework for the application of quantitative high-throughput microscopy to rapidly develop serological assays for emerging virus infections.

## 1. Introduction

The recent emergence of the novel pathogenic coronavirus SARS-CoV-2 ^[1–3]^ and the rapid pandemic spread of the virus has dramatic consequences in all affected countries. In the absence of a protective vaccine or a causative antiviral therapy for COVID-19 patients, testing for SARS-CoV-2 infection and tracking of transmission and outbreak events are of paramount importance to control viral spread and avoid the overload of healthcare systems. The sequence of the viral genome became publicly available only weeks after the initial reports on COVID-19 via the community online resource *virological*.*org* and allowed rapid development of reliable and standardized quantitative RT-PCR (qPCR) based tests for direct virus detection in nasopharyngeal swab specimens ^[4–6]^. These tests are the key to identify acutely infected individuals and monitor virus load as a basis for the implementation of quarantine measures and treatment decisions.

In response to the initial wave of COVID-19 infection many countries implemented more or less severe lockdown strategies, resulting in a gradual decrease in the rate of new infections and deaths ^[7]^. With gradual release of these lockdown strategies, monitoring and tracking of SARS-CoV-2 specific antibody levels becomes highly important. Many critical aspects of the humoral immune response against SARS-CoV-2 are currently not well understood ^[8]^. In addition, levels of infection in the general population in different areas remain largely unknown due to proportion of undocumented cases arising from asymptomatic individuals ^[9,10]^ which had not been subjected to RNA testing, or to limitations in testing capacity especially in areas of relatively high prevalence. Public health control strategies aiming at regulating human mobility and social behaviour in order to suppress the infection rate will have to take into account the proportion of seropositive individuals in the general population, or in specific population groups ^[11]^. Information on the level of antiviral antibodies, as well as on the serological response against different viral proteins, is also a key element of understanding the nature, development and durability of the antiviral immune response. Therefore, specific, sensitive and reliable methods for the quantitative detection of virus specific antibodies in human specimens are urgently needed from the beginning of an emerging pandemic.

Compared to approaches for direct virus diagnostics by PCR, development of test systems for detection of SARS-CoV-2 specific antibodies proved to be more challenging. In particular, cross reactivity of antibodies against circulating common cold coronaviruses (strains OC43, NL63, 229E and HKU1) are of concern in this respect as it was observed in case of serological tests developed for closely related SARS-CoV and MERS-CoV ^[12]^. Developments in the past months yielded well validated, commercially available ELISA or (electro)chemoluminescence-based kits for SARS-CoV-2 serological diagnostics. However, initially marketed test kits underwent a very rapid development and approval process due to the emergency of the situation, with low numbers of samples used for validation; consequently, sensitivity and specificity of the test systems often failed to meet the practical requirements ^[13]^. Furthermore, the disruption of supply chains and high demand for tests during pandemic situations can lead to shortage of commercially available test kits and/or required reagents, as witnessed in the early phases of the ongoing SARS-CoV-2 pandemic. Thus, complementary strategies to test for antiviral antibodies that can be rapidly deployed in situations where commercially available kits are either not yet developed or not available are an important addition to the diagnostic toolkit.

Immunofluorescence (IF) using virus infected cells as a specimen is a classical serological approach in virus diagnostics and has been applied to coronavirus infections, including the closely related virus SARS-CoV ^[14–16]^. The advantages of IF are (i) that it does not depend on specific diagnostic reagent kits or instruments, (ii) that the specimen contains all viral antigens expressed in the cellular context and (iii) that the method has the potential to provide high information content (differentiation of staining patterns and intensities due to reactivity against various viral proteins). A mayor disadvantage of the IF approach as it is typically used in serological testing is its limited throughput capacity due to the involvement of manual microscopy handling steps and sample evaluation based on visual inspection of micrographs. Furthermore, visual classification is subjective and thus not well standardized and yields only binary results. Here, we address those limitations, making use of advanced automated microscopy and image analysis strategies developed for basic research. We present the establishment and validation of a semi-quantitative, semi-automated workflow for SARS-CoV-2 specific antibody detection. With its 96-well format, semi-automated microscopy and automated image analysis workflow it combines advantages of IF with a reliable and objective semi-quantitative readout and high throughput compatibility. The protocol described here was developed in response to the emergence of SARS-CoV-2, but it represents a general approach that can be adapted for the study of other viral infections and is suitable for rapid deployment to support diagnostics of emerging viral infections in the future.

## 2. Results

### 2.1 Setup of the IF assay for SARS-CoV-2 antibody detection

We decided to use cells infected with SARS-CoV-2 as samples for our IF analyses, since this setup provides the best chance for detection of antibodies targeted at the different viral proteins expressed in the host cell context. African green monkey kidney epithelial cells (VeroE6 cell line) have been used for infection with SARS-CoV-2, virus production and IF^[3,17]^. In preparation for our analyses we compared different cell lines for use in infection and IF experiments, but all tested cell lines were found to be inferior to VeroE6 cells for our purposes (see Materials and Methods and Fig. S1). All following experiments were thus carried out using the VeroE6 cell line.

In order to allow for clear identification of positive reactivity in spite of a variable and sometimes high nonspecific background from human sera, our strategy involves a direct comparison of the IF signal from infected and non-infected cells in the same sample. Preferential antibody binding to infected compared to non-infected cells indicates the presence of specific SARS-CoV-2 antibodies in the examined serum. Under our conditions, infection rates of ∼40-80% of the cell population were achieved, allowing for a comparison of infected and non-infected cells in the same well of the test plate. An antibody that detects dsRNA produced during viral replication was used to distinguish infected from non-infected cells within the same field of view (Fig. 1A).

**Figure 1:**
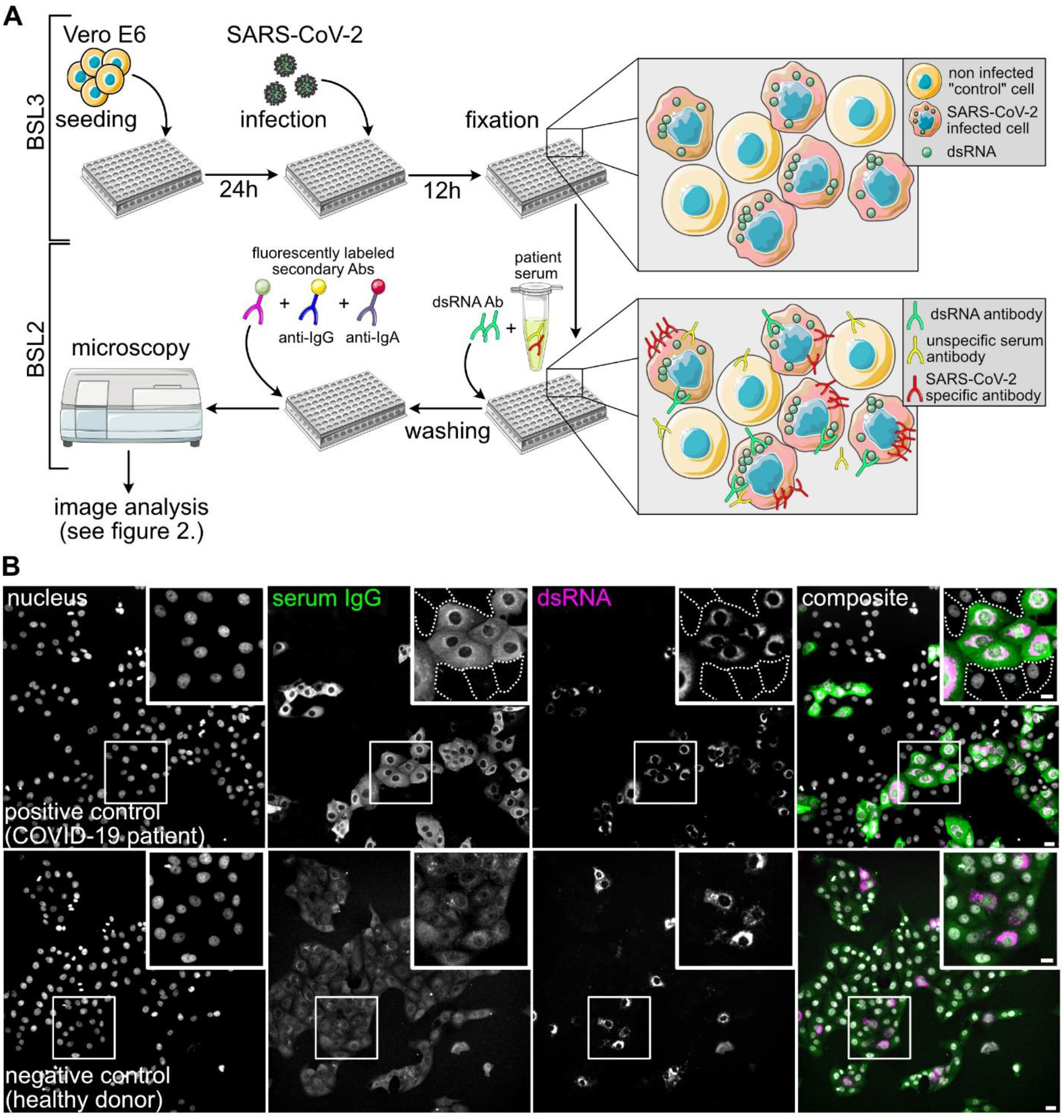
Principle of the immunofluorescence assay for SARS-CoV-2 antibody detection. (**A**) Scheme of the IF workflow and the concept for SARS-CoV-2 antibody detection. (**B**) Representative images showing immunofluorescence results using a COVID-19 patient serum (positive control, upper panels) and a negative control serum (lower panels), followed by staining with an AlexaFluor488-coupled anti-IgG secondary antibody. Nuclei (grey), IgG (green), dsRNA (magenta) channels and a composite image are shown. White boxes mark the zoomed areas. Dashed lines mark borders of non-infected cells which are not visible at the chosen contrast setting. Note that the upper and lower panels are not recorded and displayed with the same brightness and contrast settings. In the lower panels the brightness and contrast scales have been expanded in order to visualize cells in the IgG serum channel where only background staining was detected. Scale bar is 20 µm in overview and 10 µm in the insets.

In order to define the conditions for immunostaining using human serum, we selected a small panel of negative and positive control sera. Four sera from healthy donors collected before November 2019 were chosen as negative controls, and eight sera from PCR confirmed COVID-19 inpatients collected at day 14 or later post symptom onset were employed as positive controls. Sera from this test cohort were used for primary staining, and bound antibodies were detected using fluorophore-coupled secondary antibodies against human IgG, IgA or IgM.

No difference between infected and non-infected cells in serum IgG antibody binding was observed when sera collected before the onset of the SARS-CoV-2 pandemic were examined (Fig. 1B, Fig. S2). In contrast, COVID-19 patient sera were clearly characterized by higher serum IgG antibody binding to infected compared to non-infected cells (Fig. 1B). All eight COVID-19 patient serum samples yielded higher IgG binding to infected compared to non-infected cells as assessed by visual inspection (Fig. S2). Similar results were obtained when an IgA or IgM specific secondary antibody was used for detection (Fig. S3). In order to allow for the parallel assessment of IgG and IgA or IgM antibodies, we established conditions for the parallel detection of anti-IgG coupled to AlexaFluor488 and anti-IgA or anti-IgM coupled to DyLight650 or AlexaFluor647 secondary antibodies, respectively, without signal bleedthrough. Using this approach, it was possible to implement detection of SARS-CoV-2 specific IgG and IgA or IgM antibodies in a single experimental setup (Fig. S4).

Titration experiments were performed with positive control sera to determine the optimal range of serum concentration in the IF experiments. All eight positive control samples showed visually detectable specific labelling of infected cells over the range of 1:10^2^ and 1:10^5^, demonstrating robustness of the assay (Fig. S5). Serum concentrations of less than 1:10^5^ did not yield detectable signals in all cases. We decided to employ a dilution of 1:10^2^ in the further experiments

### 2.2 Image analysis

Our next aim was to establish a semi-automated analysis workflow for image acquisition and analysis for a medium to high throughput setting. VeroE6 cells were seeded into 96-well plates infected and immunostained using anti-dsRNA antibody and patient serum, followed by indirect detection using a mixture of anti-IgG and anti-IgA/IgM secondary antibodies. Images were acquired using an automated widefield microscope (see Materials and Methods section for more detail).

To obtain a measure for specific antibody binding we performed automated segmentation of cells and classified them into infected and non-infected cells based on the dsRNA staining. We then measured fluorescence intensities in the serum channel per cell as a proxy for the amount of bound antibodies for both infected and non-infected cells and calculated the ratio between these values for infected and non-infected cells in a given specimen. To enable training of a machine learning approach for cell segmentation and to directly evaluate infected cell classification, we manually labelled cells and annotated them as infected/non-infected in 10 images chosen from 5 positive and 5 control specimens. Fig. 2 presents a graphical overview of all analysis steps; the full description of every step can be found in Materials and Methods. Briefly, our approach works as follows:

**Figure 2:**
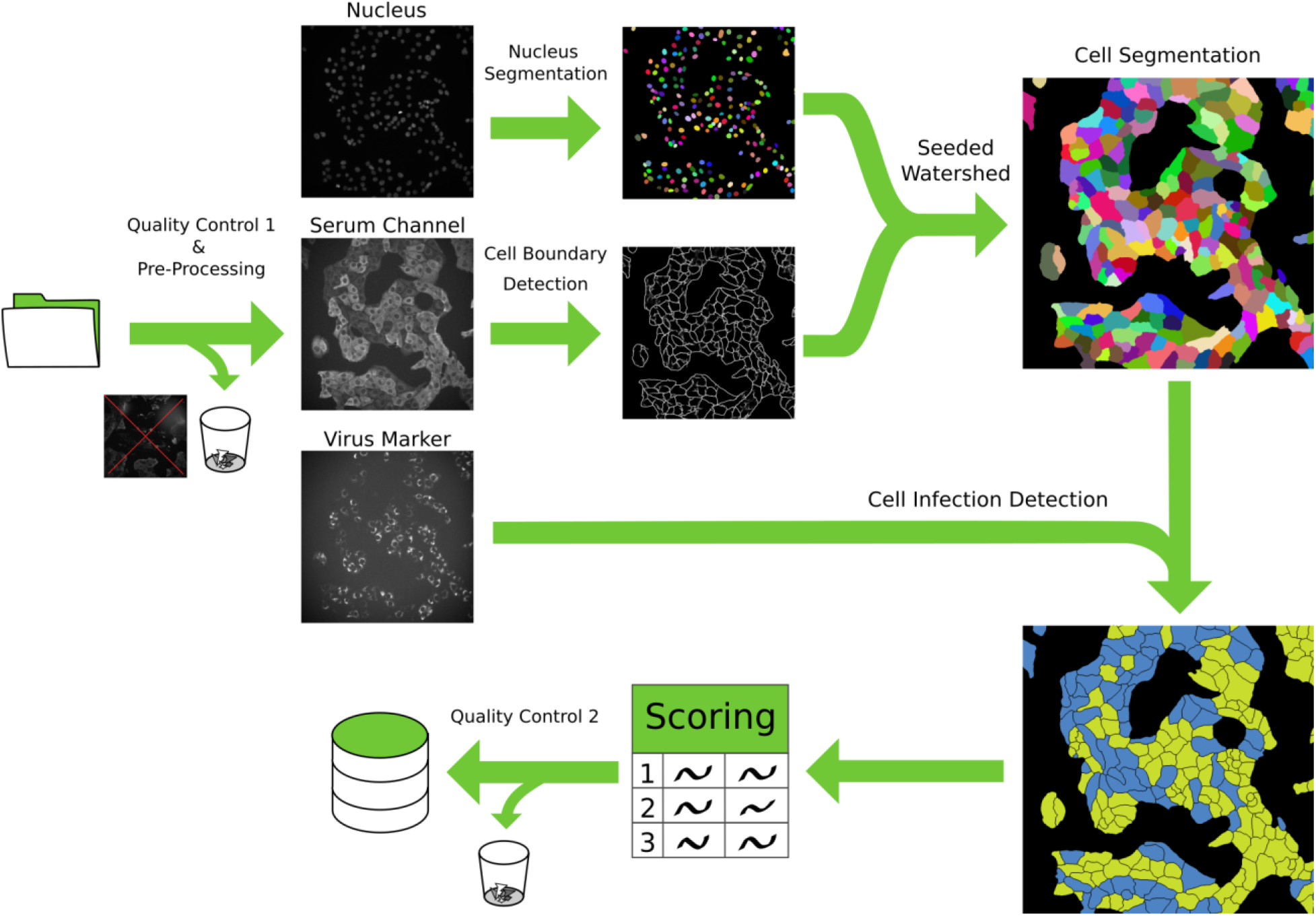
Schematic overview of the image processing pipeline. Initially, images are subjected to the first manual quality control, where images with acquisition defects are discarded. A pre-processing step is then applied to correct for barrel artifacts. Subsequently, segmentation is obtained via seeded watershed, this algorithm requires seeds obtained from StarDist segmentation of the nuclei and boundary evidence computed using a neural network. Lastly, using the virus marker channel we classify each cell as infected or not infected and we computed the scoring. A final automated quality control identifies and automatically discards non-conform results. All intermediate results are saved in a database for ensuring fully reproducibility of the results.

First, we manually discarded all images that contained obvious artefacts such as large dust particles or dirt and out-of-focus images. Then, images were processed to correct for the uneven illumination profile in each channel. Next, we segmented individual cells with a seeded watershed algorithm ^[18]^, using nuclei segmented via StarDist ^[19]^ as seeds and boundary predictions from a U-Net ^[20,21]^ as a heightmap. We evaluated this approach using leave-one-image-out cross-validation on the manual annotations and measured an average precision^[22]^ of 0.77 +-0.08 (i.e., on average 77% of segmented cells are matched correctly to the corresponding cell in the annotations). Combined with extensive automatic quality control which discards outliers in the results, the segmentation was found to be of sufficient quality for our analysis, especially since robust intensity measurements were used to reduce the effect of remaining errors.

We then classified the segmented cells into infected and non-infected, by measuring the 95th percentile intensities in the dsRNA channel and classifying cells as infected if this value exceeded 4.8 times the noise level, determined by the mean absolute deviation. This factor and the percentile were determined empirically using grid search on the manually annotated images (see above). Using leave-one-out cross validation on the image level, we found that this approach yields an average F1-score of 84.3%.

In order to make our final measurement more reliable, we then discarded whole wells, images or individual segmented cells based on quality control criteria that were determined by inspection of initial results. Those criteria include a minimal number of non-infected cells per well; minimal and maximal number of cells per image; minimal cell intensities for images; and minimal and maximal sizes of individual cells (see Materials and Methods for full details).

To score each sample, we computed the intensity ratio *r* :

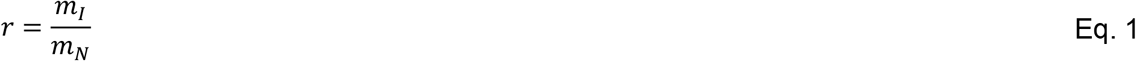

Here, *m*_*I*_ is the median serum intensity of infected cells and *m*_*N*_ the median serum intensity of non-infected cells. For each cell, we compute its intensity by computing the mean pixel intensity in the serum channel (excluding the nucleus area where we typically did not observe serum binding) and then subtracting the background intensity, which is measured on two control wells that did not contain any serum.

We used efficient implementations for all processing steps and deployed the analysis software on a computer cluster in order to enhance the speed of imaging data processing. For visual inspection, we have further developed an open-source software tool (PlateViewer) for interactive visualization of high-throughput microscopy data ^[23]^. PlateViewer was used in a final quality control step to visually inspect positive hits. For example, PlateViewer inspection allowed identifying a characteristic spotted pattern co-localizing with the dsRNA staining (Fig. S6) that was sometimes observed in the IgA channel upon staining with negative control serum. In contrast, sera from COVID-19 patients typically displayed cytosol, ER-like and plasma membrane staining patterns in this channel (Fig. 1B, Fig. S3). The dsRNA co-localizing pattern observed for sera from the negative control cohort is by definition non-specific for SARS-CoV-2, but would be classified as a positive hit based on staining intensity alone. Using PlateViewer, we performed a quality control on all IgA positive hits and removed those displaying the spotted pattern colocalising with the dsRNA signal from further analysis.

### 2.3 Assay characterization and validation

With the immunofluorescence protocol and automated image analysis in place we proceeded to test a larger number of control samples in a high throughput compatible manner for assay validation. All samples were processed for IF as described above, and in parallel analysed by a commercially available semi-quantitative SARS-CoV-2 ELISA approved for diagnostic use (Euroimmun, Lübeck, Germany) for the presence of SARS-CoV-2 specific IgG and IgA antibodies.

As outlined above, a main concern regarding serological assays for SARS-CoV-2 antibody detection is the occurrence of false positive results. A particular concern in this case is cross-reactivity of antibodies that originated from infection with any of the four types of common cold Corona viruses (ccCoV) circulating in the population. The highly immunogenic major structural proteins of SARS-CoV-2 nucleocapsid (N) and spike (S) protein, have an overall homology of ∼30% ^[3]^ to their counterparts in ccCoV and subdomains of these proteins display a higher degree homology; cross-reactivity with ccCoV has been discussed as the major reason for false positive detection in serological tests for closely related SARS-CoV and MERS-CoV ^[12]^. Also, acute infection with Epstein-Barr virus (EBV) or cytomegalovirus (CMV) may result in unspecific reactivity of human sera ^[24,25]^. We therefore selected a negative control panel consisting of 218 sera collected before the fall of 2019, comprising samples from healthy donors (n=105, cohort B), patients that tested positive for ccCoV several months before the blood sample was taken (n=34, all four types of ccCoV represented; cohort A), as well as patients with diagnosed *Mycoplasma pneumoniae* (n=22; cohort Z), EBV or CMV infection (n=57, cohort E). We further selected a panel of 57 sera from 29 RT-PCR confirmed COVID-19 patients collected at different days post symptom onset as a positive sample set (cohort C, see below).

Sera were employed as primary antisera for IF staining using IgM, IgA or IgG specific secondary antibodies, and samples were imaged and analysed as described above. This procedure yielded a ratiometric intensity score for each serum sample. Based on the scores obtained for the negative control cohort and the patient sera, we defined the threshold separating negative from positive scores for each of the antibody channels. For this, we performed ROC curve analysis ^[26–28]^ on a subset of the data (cohorts A, B, C, Z). Using this approach, it is possible to take the relative importance of sensitivity versus specificity as well as seroprevalence in the population (if known) into account for optimal threshold definition. By giving more weight to false positive or false negative results, one can adjust the threshold dependent on the context of the study. Whereas high sensitivity is of importance for e.g. monitoring seroconversion of a patient known to be infected, high specificity is crucial for population based screening approaches, where large study cohorts characterized by low seroprevalence are tested. Since we envision the use of the assay for screening approaches, we decided to assign more weight to specificity at the cost of sensitivity for our analyses (see Materials and Methods for an in-depth description of the analysis). Optimal separation in this case was given using threshold values of 1.39, 1.31 and 1.27 for IgA, IgG and IgM channels respectively (Fig. S7). We validated the classification performance on negative control cohort E (n=57) which was not seen during threshold selection, and detected no positive scores. Results from the analysis of the negative control sera are presented in Fig. 4 and Table 1.

**Table 1:** Summary of positive results for the negative control samples obtained by ELISA and IF. The classification of positive or borderline results in ELISA followed the definition of the test manufacturer. The classification in IF is described in materials and methods. Positive IgA and IgG ELISA readings were derived from the same sample. Cohort B = healthy donors, cohort A = patients that tested positive for ccCoV (all four types of ccCoV represented), cohort Z = patients with diagnosed *Mycoplasma pneumoniae*, cohort E = patient with diagnosed EBV or CMV infection. ^a^ – borderline values were considered positive.

| Negative cohort | IF IgM | IF IgA | IF IgG | ELISA IgA | ELISA IgG |
| --- | --- | --- | --- | --- | --- |
| **B (n=105)** | 1 | 0 | 1 | 7 | 5 |
| **A (n=34)** | 0 | 0 | 1 | 3 | 1 |
| **Z (n=22)** | 0 | 0 | 0 | 2 | 0 |
| **E (n=57)** | 0 | 0 | 0 | 11 | 1 |
| **Total (n=218)** | **1 (0,5%)** | **0 (0,0%)** | **2 (0,9%)** | **23<sup>a</sup> (10,6%)** | **7<sup>a</sup> (3,2%)** |

**Figure 3:**
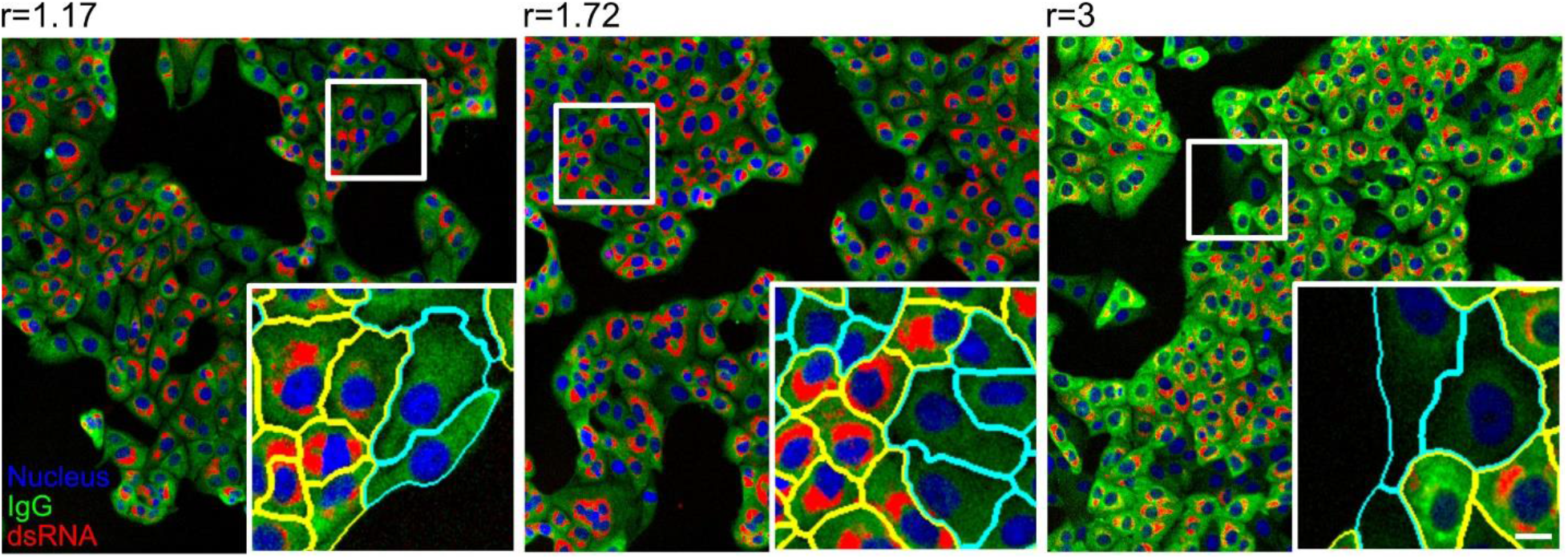
Examples of results from the automated image analysis pipeline. Panels display images that correspond to three different ratio scores (ratio score is indicated above the image) determined from samples stained with three different human sera, followed by staining with an anit-IgG secondary antibody coupled to AlexaFluore488. Images represent overlays of three channels - nuclei (blue), IgG (green) and dsRNA (red). White boxes mark the zoomed area. Cells in the insets are highlighted with yellow or cyan boundaries, indicating infected and non-infected cells, respectively. Scale bar = 10 μm.

**Figure 4:**
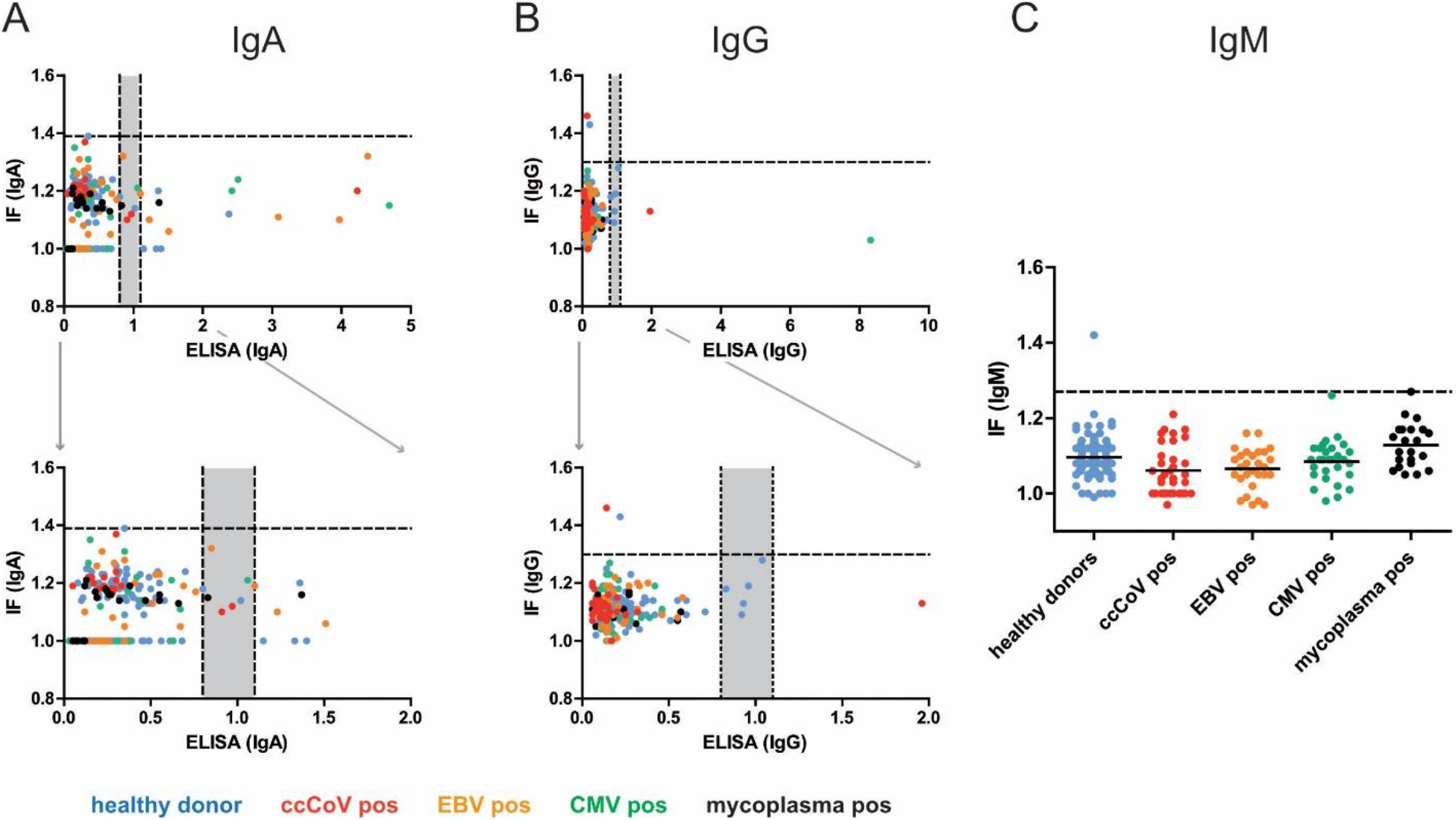
Correlation between SARS-CoV-2 specific IF and ELISA results for the negative control panel obtained in IgA (A) or IgG (B) measurements. Each dot represents one serum sample. Blue, healthy donors; red, ccCoV positive; green, CMV positive; orange, EBV positive; black, mycoplasma positive. Bottom panels represent zoomed-in versions of the respective top panel to illustrate the borderline region. **(C)** IgM values for the indicated negative control cohorts determined by IF. Since a corresponding IgM specific ELISA kit from Euroimmun was not available, correlation was not analysed in this case. In some cases, antibody binding above background was undetectable by IF in non-infected as well as in infected cells, indicating low unspecific cross-reactivity and lack of specific reactivity of the respective serum. In order to allow for inclusion of these data points in the graph, the IF ratio was set to 1,0. Dotted lines indicate the optimal separation cut-off values defined for sample classification, grey areas indicate borderline results in ELISA.

While the majority of these samples tested negative in ELISA measurements as well as in the IF analyses, some positive readings were obtained in each of the assays, in particular in the IgA specific analyses (Fig. 4 and Table 1). Since samples from these cohorts were collected between 2015 and 2019, and donors were therefore not exposed to SARS-CoV-2 before sampling, these readings represent false positives. Of note, negative control cohort E displayed a particularly high rate of false positives in ELISA measurements, but not in IF (Table 1). We conclude that the threshold values determined achieve our goal of yielding highly specific IF results (at the cost of sub-maximal sensitivity).

Roughly 10.6% (IgA) or 3% (IgG) of the samples were classified as positive or potentially positive by ELISA (Table 1). The notably lower specificity of the IgA determination in a seronegative cohort observed here is in accordance with findings in other studies ^[29,30]^ and information provided by the manufacturer of the test (90,5% for IgA vs. 99,3% for IgG; Euroimmun SARS-CoV-2 data sheet, April 24, 2020; in response to these findings, an improved version of the test has been recently developed). The respective proportion of false-positives obtained based on IF, 0% for IgA and 0,9% for IgG, were lower, indicating higher specificity of the IF readout compared to the ELISA measurements. Importantly, however, false positive readings did not correlate between ELISA and IF (Fig. 4). Thus, classifying only samples that test positive in both assays as true positives resulted in the elimination of false positive results (0 of 218 positives detected). We conclude that applying both methods in parallel and using the ‘double positive’ definition for classification notably improves specificity of SARS-CoV-2 antibody detection.

In order to determine the sensitivity of our IF assay, we employed 57 sera from 29 symptomatic COVID-19 patients that had been RT-PCR confirmed for SARS-CoV-2 infection. Archived sera from these patients had been collected in the range between day 5 and 27 post symptom onset. Again, samples were measured both in IF and ELISA, and the correlation between the semi-quantitative values was assessed as shown in Fig. 5. While there were deviations in the height of the values, positive correlation was evident in both cases, with values for the IgG readout being more congruent than those for the less specific IgA determination (Pearson r: 0,847 for IgG; 0,655 for IgA).

**Figure 5:**
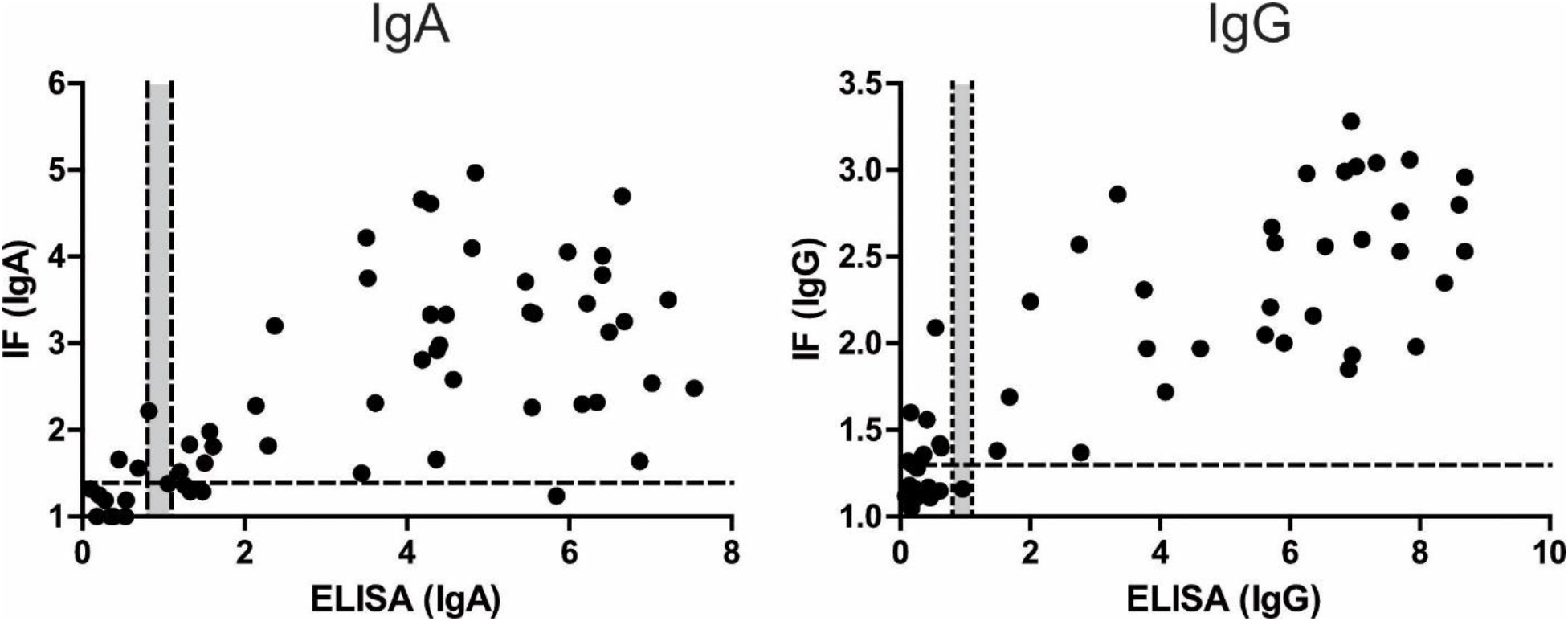
Correlation between IgA or IgG values obtained by ELISA and IF for sera from 29 COVID-19 patients collected at different days post infection. In some cases, antibody binding above background was undetectable by IF in non-infected as well as in infected cells, indicating low unspecific cross-reactivity and lack of specific reactivity of the respective serum. In order to allow for inclusion of these data points in the graph, the IF ratio was set to 1,0. Dotted lines indicate the cut-off values defined for classification of readouts, grey areas indicate borderline values.

For an assessment of sensitivity, we stratified the samples according to the day post symptom onset, as shown in Fig. 6. and Table 2. For both methods, and for all antibody classes, mean values and the proportion of positive samples increased over time. In all cases, only positive values were obtained for samples collected later than day 14 post symptom onset, in accordance with other reports ^[30–32]^. Consistent with other reports ^[32]^, SARS-CoV-2 specific IgM was not detected notably earlier than the two other antibody classes in our measurements. At the earlier time points (up to day 14), a similar or higher proportion of positive samples was detected by IF compared to ELISA for IgG. Although the sample size used here is too small to allow a firm conclusion, these results suggest that the sensitivity of IgG detection by the semi-quantitative IF approach is higher than that of an approved semi-quantitative ELISA assay routinely used in diagnostic labs. In the case of IgA detection at earlier time points (< day 11) ELISA performed slightly better (11/17 samples scored positive) compared to IF (9/17 scored positive) however that came with the price of a very low specificity of ELISA IgA assay (10.6% false negative detection) compared to IF (0.5%).

**Table 2:**
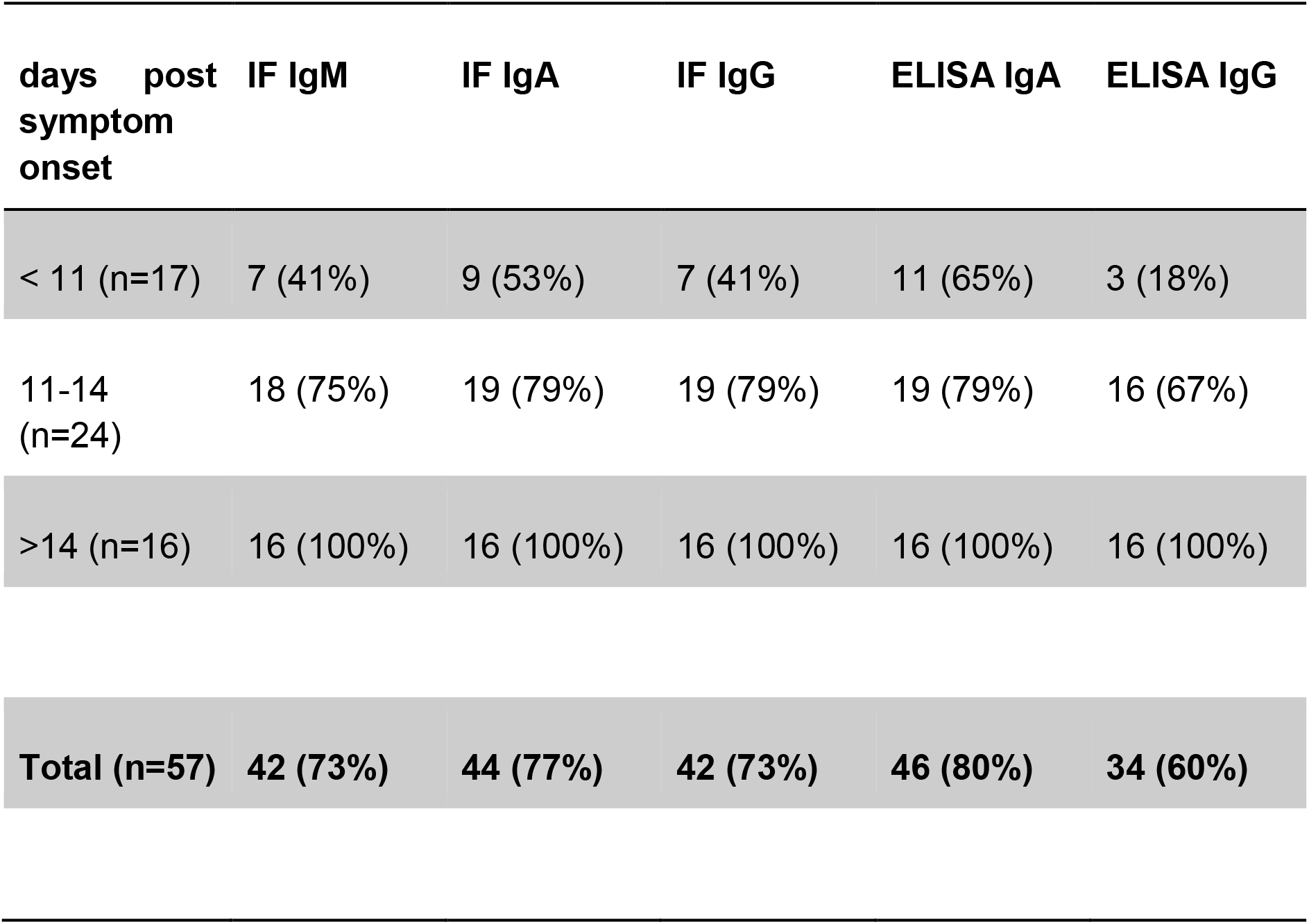
Positive results obtained for sera from COVID-19 patients collected at the indicated days post symptom onset.

| **days post symptom onset** | **IF IgM** | **IF IgA** | **IF IgG** | **ELISA IgA** | **ELISA IgG** |
| --- | --- | --- | --- | --- | --- |
| < 11 (n=17) | 7 (41%) | 9 (53%) | 7 (41%) | 11 (65%) | 3 (18%) |
| 11-14 (n=24) | 18 (75%) | 19 (79%) | 19 (79%) | 19 (79%) | 16 (67%) |
| >14 (n=16) | 16 (100%) | 16 (100%) | 16 (100%) | 16 (100%) | 16 (100%) |
| **Total (n=57)** | **42 (73%)** | **44 (77%)** | **42 (73%)** | **46 (80%)** | **34 (60%)** |

**Figure 6:**
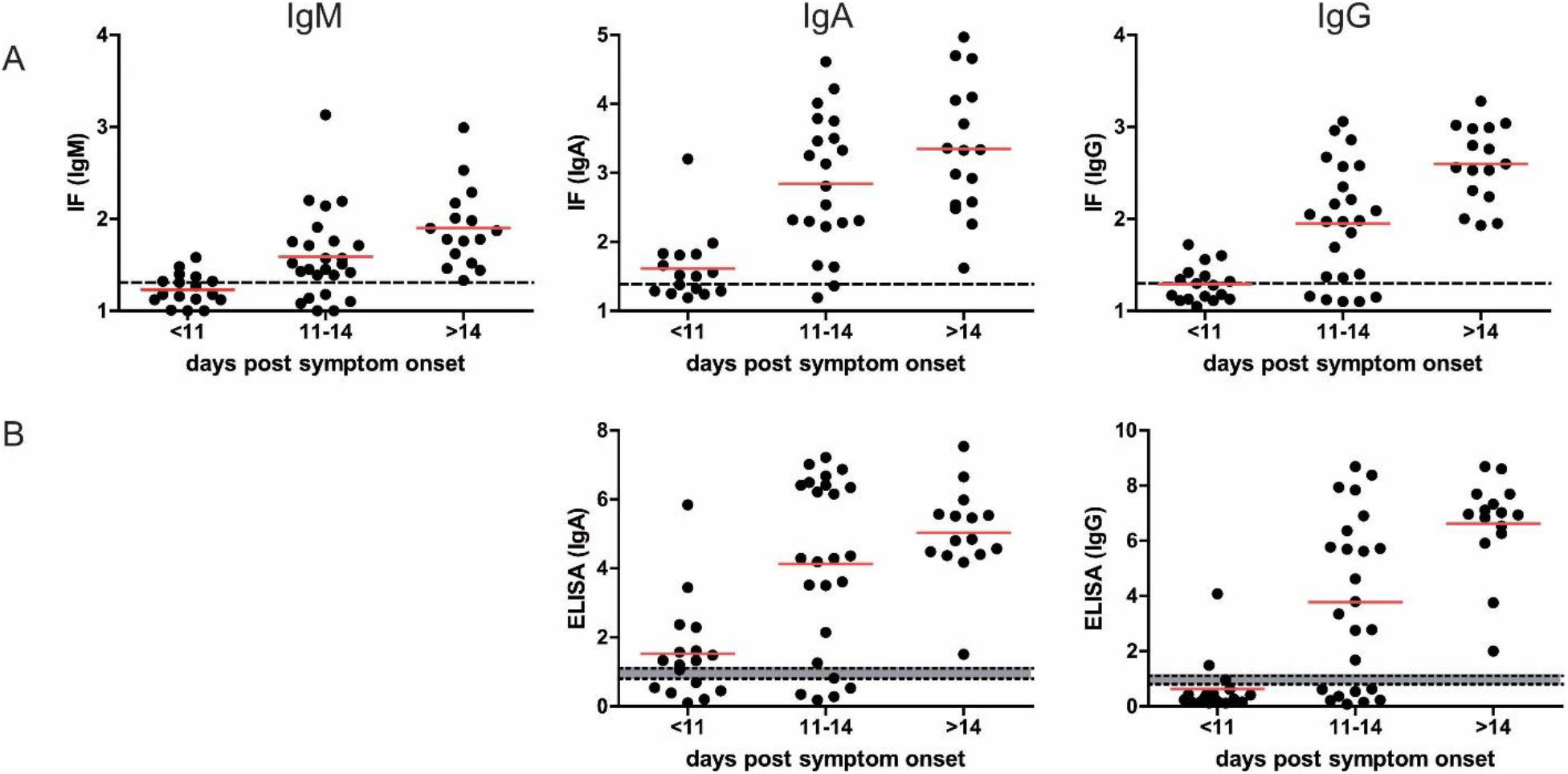
Detection of SARS-CoV-2 specific antibodies in sera from COVID-19 patients. (**A**) Fifty-seven serum samples from 29 PCR confirmed patients collected at the indicated times post symptom onset were analysed by the IF workflow for the presence of SARS-CoV-2 specific IgM, IgA and IgG antibodies. Each dot represents one serum sample. Red line: mean value; dotted line: cut-off between negative and positive values. (**B**) The same samples as in A were analysed by ELISA for the presence of SARS-CoV-2 specific IgA and IgG antibodies. Each dot represents one serum sample. Red line: mean value; dotted lines: cut-off; grey zone: borderline.

## 3. Discussion

Here, we describe the development of a semi-quantitative IF based assay for detection of SARS-CoV-2 specific antibodies in human samples that complements available ELISA-based testing systems ^[33,34]^. Alternatives to ELISA-based commercial test kits are important in situations where those kits are not available either because they are not yet developed in early days of the pandemic or due to high global demands for tests and required reagents. The microscopy-based assay described here has been developed during the early phase of the COVID-19 pandemic to support the serological testing needs of the University Hospital Heidelberg, Germany and is employed as a confirmatory assay in clinical studies ^[35]^ and ongoing studies]. The assay displayed comparable or slightly better sensitivity and specificity than a commercially available semi-quantitative SARS-CoV-2 ELISA approved for diagnostic use at the time. More importantly, combining two technically different serological assays, IF and ELISA, and classifying as “positive hits” only those that scored positive in both assays was instrumental to minimize false positive results while maintaining high sensitivity, and thus serves as a principle for serological studies or diagnostics where specificity of detection is of critical importance. Specificity of detection is essential in settings of relatively low SARS-CoV-2 antibody prevalence ^[36–38]^ in conjunction with high prevalence of potentially cross-reactive anti-ccCoV antibodies in a global population ^[39]^.

One advantage of the IF based assay presented here is that the specimens used for detection present the entire viral proteome, while ELISA or chemiluminescent approaches use a single recombinantly expressed antigen. Both the N and S protein of coronaviruses are highly immunogenic, and antibodies binding to the receptor binding domain on the S1 subunit are considered most relevant for neutralization. However, the relative importance of antibodies directed against the N protein for potential protective immunity against SARS-CoV-2 and the possible relevance of the overall breadth of the antibody response is currently unclear. Other SARS-CoV-2 structural and non-structural proteins might play a role in immune response as it was shown for proteins 3a and 9b of the closely related SARS-CoV ^[40]^. In addition, expression of the viral proteome in permissive cells ensures correct protein folding and post-translational modification patterns. Alterations in post-translational modifications are likely to influence the ability of serum antibodies to bind to different viral epitopes as it was shown for other viruses such as HIV-1 ^[41]^. It has to be noted that the detection of viral RNA requires fixation and permeabilization of cells, which has the potential to affect epitope preservation. However, based on the high sensitivity of antibody detection and the good correlation to ELISA measurements observed we conclude that this was no major concern in this case.

Two major disadvantages of typical IF-based serological assays as applied in the past are manual microscopy acquisition steps and evaluation of samples based on a visual inspection. This procedure is incompatible with high throughput approaches and results are subjective, not quantitative and difficult to standardize. We have addressed these disadvantages by implementing automated microscopy acquisition and developing a robust software platform that is able to identify individual cells, classify infected and non-infected cells and take into account specific and non-specific background in order to generate semi-quantitative results. Depending on the context of a study and the questions to be addressed, sensitivity or specificity may be of higher importance. The automated image analysis protocol developed here allows the user to adapt the classification according to the study needs, putting more weight on either one of the parameters.

Automated image acquisition and image analysis presented here are compatible with a high throughput approach. Plates with fixed samples of infected cells can be prepared in advance and stored at 4°C for several weeks. In the manual workflow used here, four 96-well plates (384 samples) could easily be analysed within a typical work day (1.5 h for immunofluorescence, 1.5 h for image acquisition, 2 h of image analysis). This is already the throughput in the range of some ELISA-based automated systems used in diagnostics and is sufficient for urgent applications in an early phase of disease response. The major disadvantage of the procedure described here for a virus like SARS-CoV-2 is the requirement of a BSL3 containment area to generate virus stocks and produce infected cell specimens. Recombinant cell lines expressing key viral antigens can address this drawback and also allows to easily implement already established automated cell seeding and immunostaining pipelines for a true high-throughput application ^[42,43]^. Combining such cell lines with spectral unmixing microscopy ^[44]^ would not only enable simultaneous determination of levels of all three major classes of antibodies (IgM, IgG and IgA), but also identification of the viral antigens recognized, in a single multiplexed approach. The high information content of the IF data (differential staining patterns) together with a machine learning-based approach ^[45]^ and the implementation of stable cell lines expressing selected viral antigens in the IF assay will provide additional parameters for classification of patient sera and further improve sensitivity and specificity of the presented IF assay.

The described analysis pipeline can be readily applied for serological analysis of other virus infections, provided that an infectable cell line and a staining procedure that allows differentiating between infected and non-infected cells are available. The assay described here thus offers potential as an immediate response to any future virus pandemic, as it can be rapidly deployed from the moment the first isolate of the pathogen has been obtained without requiring information on the expression of immunogenicity of viral proteins.

## 4. Materials and Methods

### 4.1 Human material

Negative control serum samples (n=218) were collected for various serological testing in the routine laboratory of the Center of Infectious Diseases, University Hospital Heidelberg between 2015 and 2019, before the start of the SARS-CoV-2 outbreak. Samples used corresponded to pseudonymized remaining material from the archive of the Center of Infectious Diseases Heidelberg. SARS-CoV-2 positive sera were collected from 29 PCR confirmed symptomatic COVID-19 inpatients (n=17) or outpatients (n=12) treated at the University Hospital Heidelberg under general informed consent (ethics votum no S-148/2020, University Hospital Heidelberg). Days post symptom onset were defined based on the anamnesis carried out upon admission. Serum samples were stored at −20°C until use.

### 4.2 Virus stock production

VeroE6 cells were cultured in Dulbecco’s modified Eagle medium (DMEM, Life Technologies) containing 10% fetal bovine serum, 100 U/mL penicillin, 100 µg/mL streptomycin and 1% non-essential amino acids (complete medium).

SARS-CoV-2 virus stocks were produced by amplification of the BavPat1/2020 strain (European Virus Archive) in VeroE6 cells. To generate the seed virus (passage 3), VeroE6 cells were infected with the original virus isolate, received as passage 2, at an MOI of 0.01. At 48 h post infection (p.i.), the supernatant was harvested and cell debris was removed by centrifugation at 800xg for 10 min. For production of virus stocks (passage 4), 500µl of the seed virus was used to infect 9×10^6^ VeroE6 cells. The resulting supernatant was harvested 48h later as described above. Virus titers were determined by plaque assay. Briefly, 2.5×10^6^ VeroE6 cells were plated into 24 well plates. 24 h later, cells were infected with serial dilutions of SARS-CoV-2 for 1 h. Inoculum was then removed and the cells were overlaid with serum free DMEM containing 0.8% carboxymethylcellulose. At 72 h. p.i., cells were fixed with 5% formaldehyde for 1 h followed by staining with 1% Crystal violet solution. Plaque forming units per ml (PFU/ml) were estimated by manual counting of the viral plaques. Stock solutions were stored in aliquots at −80°C until use for infection experiments.

### 4.3 Infection of cells and immunofluorescence staining

In order to find a suitable cell line for our application, we performed pre-experiments comparing different cell lines with respect to their susceptibility to SARS-CoV-2 infection. Cells were seeded on glass coverslips and infected on the following day with SARS-CoV-2 strain BavPat1/2020 for 16h at MOI 0.01. Cells were fixed with 6%PFA in PBS, followed by permeabilisation with 0.5% Triton X100 in PBS and then subjected to a standard immunofluorescence staining protocol as described in materials and methods. Only very few infected calls were detected in the case of hepatocyte-derived carcinoma cells (HUH-7), human embryonic kidney (HEK293T) and human alveolar basal epithelial (A549) cells (Fig. S1). Calu-3 cells grew in small clumps, often on top of each other which impacted our microscopy-based readout. In contrast, VeroE6 cells grew as a monolayer and were viable for at least 24 h p.i. Based on these results, VeroE6 cells were chosen for all experiments in this manuscript.

For serum screening by IF microscopy, VeroE6 cells were seeded at a density of 7,000 cells per well into a black-wall glass-bottom 96 well plates (Corning, Product Number 353219) or on glass coverslips placed in a 24-well plate. 24 h after seeding, cells were infected with SARS-CoV-2 at an MOI of 0.01 for 16 h. Cells were then fixed with 6% Formaldehyde for 1 h followed by washing 3x with phosphate buffered saline (PBS) under biosafety level 3. Afterwards, samples were handled under biosafety level 2. Cells were washed once in PBS containing 0,02% Tween 20 (Sigma) and permeabilised using 0,5% Triton X100 (Sigma) for 10 minutes. Samples were washed again and blocked using 2% powdered milk (Roth) in PBS for 20 min followed by two additional washing steps. All washing steps in a 96-well format were performed using the HydroFlex microplate washer (Tecan). Next, cells were incubated with patient serum (prediluted 1:1 in 0,4% Triton-X100 in PBS; further dilution 1:50 in PBS if not stated otherwise) and anti-ds-RNA mouse monoclonal J2 antibody (Scicons, 1:4000) in PBS for 30 min at room temperature. After 3 washing steps, cognate secondary antibodies were applied for 20 min at room temperature. Goat anti-human IgG-AlexaFluor 488 (Invitrogen, Thermofisher Scientific), goat anti-human IgA DyLight 650 (Abcam), goat anti-human IgM u chain (Invitrogen, Thermofisher Scientific), for detecting immunoglobulins in human serum together with goat anti-mouse IgG-AlexaFluor 568 (Invitrogen, Thermofisher Scientific) for dsRNA detection, all at 1:2000 dilution in PBS, have been used. After incubation with secondary antibodies cells were washed twice, stained with Hoechst (0,002µg/ml in PBS) for 3 minutes, washed again twice and stored at +4°C until imaging.

### 4.4 Microscopy

Samples were imaged on motorized Nikon Ti2 widefield microscope using a Plan Apo lambda 20x/0.75 air objective and a back-illuminated EM-CCD camera (Andor iXon DU-888). To automatically acquire images in 96-well format, the JOBS module was used. The system was configured to acquire 9 images per well (in a regular 3 x 3 pattern centered in the middle of each well). The Perfect Focus System was used for autofocusing followed by a software-based fine focusing using the Hoechst signal in an axial range of 40um. Images were acquired in 4 channels using the following excitation/emission settings: Ex 377/50, Em 447/60 (Hoechst); Ex 482/35, Em 536/40 (AlexaFluor 488); Ex 562/40, Em 624/40 (AlexaFluor 568) and Ex 628/40, Em 692/40 (AlexaFluor 647 and DyLight 650). Exposure times were in the range between 50 and 100ms with EM gain between 50 and 150.

### 4.5 Enzyme linked immuno-sorbent assay (ELISA)

ELISA measurements for determination of reactivity against the S1 domain of the viral spike protein were carried out using the Euroimmun Anti-SARS-CoV-2-ELISA (IgA) and Anti-SARS-CoV-2-ELISA (IgG) test kits (Euroimmun, Lübeck, Germany; EI 2606-9601 A and EI 2606-9601 G) run on an Euroimmun Analyzer I instrument according to the manufacturer’s instructions. Optical densities measured for the samples were normalized using the value obtained for a calibrator sample provided in the test kit. The interpretation of the semi-quantitative ratiometric values obtained followed the manufacturer’s protocol: values <0.8 were classified as negative, 0.8-1.1 as borderline, and values of 1.1 or higher as positive.

### 4.6 Image Analysis

#### Manual Annotations

Two of our processing steps require manually annotated data: in order to train the convolutional neural network used for boundary and foreground prediction, we needed label masks for the individual cells. To determine suitable parameters for the infected cell classification, we needed a set of cells classified as being infected or non-infected. We have produced these annotations for 10 images with the following steps. First, we created an initial segmentation following the approach outlined in the Segmentation subsection, using boundary and foreground predictions from the ilastik ^[46]^ pixel classification workflow, which can be obtained from a few sparse annotations. We then corrected this segmentation using the annotation tool BigCat (https://github.com/saalfeldlab/bigcat). After correction, we manually annotated these cells as infected or non-infected. Note that this mode of annotations can introduce two types of bias: the segmentation labels are derived from an initial segmentation. Small systematic errors in the initial segmentation that were not found during correction, could influence the boundary prediction network. More importantly, when annotating the infected / non-infected cells, both the serum channel and the virus marker channel have to be available to the annotators, in order to visually delineate the cells. This may result in subconscious bias, with the observed intensity in the serum channel influencing the decision on the infection status of a cell.

#### Preprocessing

On all acquired images, we performed minimal preprocessing (i.e., flat-field correction) in order compensate for uneven illumination of the microscope system ^[47]^. First, we subtract a constant CCD camera offset (ccd_offset). Secondly, we correct uneven illumination by dividing each channel by a corresponding corrector image (flatfield(*x, y*)), which was obtained as a normalized average of all images of that channel, smoothed by a normalized convolution with a Gaussian filter with a bandwidth of 30 pixels.

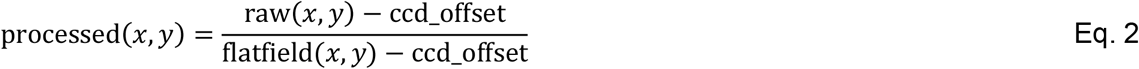

This corrector image was obtained for all images of a given microscope set-up. Full background subtraction is performed later in the pipeline using either the background measured on wells that (deliberately) do not contain any serum or, if not available, using a fixed value that was determined manually.

#### Segmentation

Cell segmentation forms the basis of our analysis method. In order to obtain an accurate segmentation, we make use of both the DAPI and the serum channel. First, we segment the nuclei on the DAPI channel using the StarDist method ^[19]^ trained on data from Caicedo et al. 2019 ^[48]^. Note that this method yields an *instance segmentation*: each nucleus in the image is assigned a unique ID. In addition, we predict per pixel probabilities for the boundaries between cells and for the foreground (i.e. whether a given pixel is part of a cell) using a 2D U-Net ^[20]^ based on the implementation of Wolny et al. 2020 ^[21]^. This method was trained using the 9 annotated images, see above. The cells are then segmented by the seeded watershed algorithm ^[18]^. We use the nucleus segmentation, dilated by 3 pixels, as seeds and the boundary predictions as the height map. In addition, we threshold the foreground predictions, erode the resulting binary image by 20 pixels and intersect it with the binarised seeds. The result is used as a foreground mask for the watershed. The dilation / erosion is performed to alleviate issues with very small nucleus segments / imprecise foreground predictions. In order to evaluate this segmentation method, we train 9 different networks using leave-one-out cross-validation, training each network on 8 of the manually annotated images and evaluating it on the remaining one. We measure the segmentation quality using average precision ^[22]^ at an intersection over union (IoU) threshold of 0.5 as described in https://www.kaggle.com/c/data-science-bowl-2018/overview/evaluation. We measure a value of 0.77 +-0.08 with the optimum value being 1.0.

#### Quantitation and Scoring

##### Infection classification

To distinguish infected cells from control cells we use the dsRNA virus marker channel: infected cells show a signal in this channel while the non-infected control cells should ideally be invisible (see Fig. 3). We classified each cell in the cell segmentation (see above) individually, using the following procedure. First, we denoised the marker channel using a white tophat filter with a radius of 20 pixels. To account for inaccuracies in the cell segmentation (the exact position of cell borders is not always clear), we then eroded all cell masks with a radius of 5 pixels and thereby discard pixels close to segment boundaries. This step does not lead to information loss, since the virus marker is mostly concentrated around the nuclei. On the remaining pixels of each cell, we compute the 0.95 quantile (*q*) of the intensity in the marker channel. For the pixels that the neural network predicts to belong to the background (*b*), we compute the median intensity of the virus marker channel across all images in the current plate. Finally, we classify the cell as infected if the 0.95 quantile of its intensity exceeds the median background by more than a given threshold:

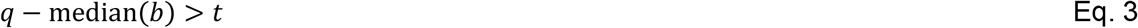

For additional robustness against intensity variations we adapt the threshold based on the variation in the background in the plate. Hence, we define it as a multiple of the mean absolute deviation of all background pixels of that plate with N=4.8:

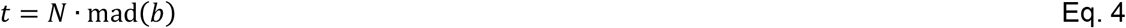

To determine the optimal values of the parameters used in our procedure, we used the cells manually annotated as infected / non-infected (see above). We performed grid search over the following parameter ranges:

- Quantile: 0.9, 0.93, 0.95, 0.96, 0.97, 0.98, 0.99, 0.995
- *N*: 0 to 10 in intervals of 0.1

To estimate the validation accuracy, we performed leave-one-out cross-validation on the image level. This yields an average validation F1 score of 84.3%, precision of 84.3% and recall of 84.8%. These values are the arithmetic means of the individual results per split.

#### Immunoglobulin intensity measurements

In order to obtain a relative measure of antibody binding, we determined the mean intensity and the integrated intensity in each segmented cell from images recorded in the IgG, IgA or IgM channel. A comparative analysis revealed that the mean intensity was more robust against the variability of cell sizes, whereas using the integrated intensity as a proxy yielded a higher variance in non-infected cells. Thus, mean intensity per cell was chosen as a proxy for the amount of antibody bound. Non-specific auto-fluorescence signals required a background correction of the measured average serum channel intensities. For background normalization, we used cells (one well per plate) which were not immunostained with primary antiserum. From this we computed the background to be the median serum intensity of all pixels of images taken from this well. This value was subtracted from all images recorded from the respective plate. In case this control well was not available, background was subtracted manually by selecting the area outside of cells in randomly selected wells and measuring the median intensity.

#### Scoring

The core interest of the assay is to measure the difference of antibody binding to cells infected with the coronavirus in comparison to non-infected control cells. To this end, utilizing the results of the image analysis, we compute the following summary statistics of the background corrected antibody binding of infected cells, *I*, and of non-infected cells, *N*:

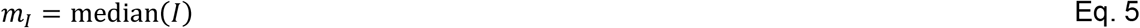

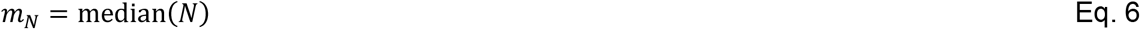

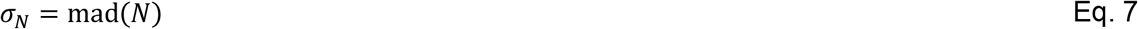

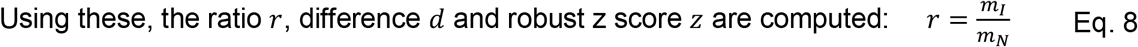

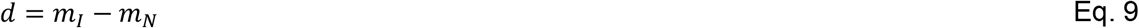

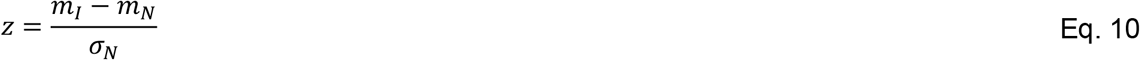

We compute above scores for each well and each image, taking into account only the cells that passed all quality control criteria (see below). While the final readout of the assay is well-based, image scores are useful for quality control.

#### Decision threshold selection

In order to determine the presence of SARS-CoV-2 specific antibodies in patient sera, it was necessary to define a decision threshold r*. If a measured intensity ratio r is above a decision threshold r* than the serum would be characterized as positive for SARS-CoV-2 antibodies. For this an ROC analysis was performed ^[28]^. Each possible choice of r* for a test corresponds to a particular sensitivity/specificity pair. By continuously varying the decision threshold, we measured all possible sensitivity/specificity pairs, known as ROC curves (Fig. S7). To determine the appropriate r* we considered two factors ^[26]^:

- The undesirability of errors or relative cost of false-positive and false-negative classifications
- The prevalence, or prior probability of disease

These factors can be combined to calculate a slope in the ROC plot^[26–28]^

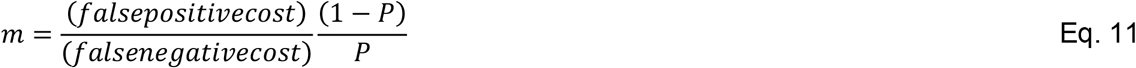

where *P* is the prevalence or prior probability of disease.

The optimal decision threshold r*, given the false-positive/false-negative cost ratio and prevalence, is the point on the ROC curve where a line with slope m touches the curve. As discussed in the main text, a major concern regarding serological assays for SARS-CoV-2 antibody detection is the occurrence of false-positive results. Therefore, we choose m to be larger than one in our analysis. In particular, we determine r* for the choice of m=10 (see Fig. S7).

#### Quality control

We performed quality control of the images and analysis results at the level of wells, images and cells. The entities that did not pass quality control are not taken into account when computing the score during final analysis. We exclude wells that contain less than 100 non-infected cells, that have a median serum intensity of infected cells smaller than 3 times the noise level (measured by the median absolute deviation), or that have negative intensity ratios, which can happen due to the background subtraction. Out of 1.736 wells, 94 did not pass the quality control, corresponding to 5.4 % of wells. At the image level, we visually inspect all images and mark those that contain imaging artifacts using a viewer based on napari ^[49]^. We distinguish the following types of artifacts during the visual inspection: empty, unstained or over-saturated images, as well as images covered by a large bright object. In addition, we automatically exclude images that contain less than 10 or more than 1000 cells. These thresholds are motivated by the observation that too few or too many cells often result from a problem in the assay. Thus, 296 of the total 15.624 images were excluded from further analysis, corresponding to 1.9 % of images. Out of these, 295 were manually marked as outliers and only a single one did not pass the subsequent automatic quality control. Finally, we automatically exclude segmented cells with a size smaller than 250 pixels or larger than 12.500 pixels that most likely correspond to segmentation errors. These limits were derived by the histogram of cell sizes investigated for several plates. Two percent of the approximate 5.5 million segmented cells did not pass this quality control. In addition, we have also inspected all samples scored as positives. For the IgA channel, we have found a dotty staining pattern in ten cases that produced positive hits based on intensity ratio in negative control cohorts, but does not appear to indicate a specific antibody response. We have also excluded these samples from further analysis.

#### Implementation

In order to scale the analysis workflow to the large number of images produced by the assay, we implemented an open-source python library to run the individual analysis steps. This library allows rerunning experiments for a given plate for newly added data on demand and caches intermediate results in order to rerun the analysis from checkpoints in case of errors in one of the processing steps. To this end, we use a file layout based on hdf5 ^[50]^ to store multi-resolution image data and tabular data. The processing steps are parallelized over the images of a plate if possible. We use efficient implementations for the U-Net ^[21]^, StarDist ^[19]^ and the watershed algorithm (http://ukoethe.github.io/vigra/) as well as other image processing algorithms ^[51]^. We use pytorch (https://pytorch.org/) to implement GPU-accelerated cell feature extraction. The total processing time for a plate (containing around 800 images) is about two hours and thirty minutes using a single GPU and 8 CPU cores. In addition, the results of the analysis as well as meta-data associated with individual plates are automatically saved in a centralized MongoDB database (https://www.mongodb.com) at the end of the workflow execution. Apart from keeping track of the analysis outcome and meta-data, a user can save additional information about a given plate/well/image in the database conveniently using the PlateViewer (see below). All source code is available open source under the permissive MIT license at https://github.com/hci-unihd/batchlib.

#### Data visualization

In order to explore the numerical results of our analysis together with the underlying image data we further developed a Fiji ^[52]^ based open-source software tool for interactive visualization of high-throughput microscopy data ^[23]^. The PlateViewer links interactive results tables and configurable scatter plots (image and well based) with a plate view of all raw, processed and segmentation images. The PlateViewer is connected to the centralised database such that also image and well based metadata can be accessed. The viewer thus enables efficient visual inspection and scientific exploration of all relevant data of the presented assay.

#### Data availability

The data from the IF assay are available in the BioImage Archive (http://www.ebi.ac.uk/bioimage-archive) under accession number S-BIAD24. This includes raw microscopy images, intermediate segmentation and infected cell classification results as well as quality control and final score results.

## Acknowledgements

We would like to thank Martin Weigert and Uwe Schmidt for their help with setting up prediction for StarDist. We would like to acknowledge Infectious Disease Imaging Platform (IDIP) at Center for Integrative Infectious Diseases Research (CIID) for microscopy support. We would like to thank EMBL, especially the EMBL IT Services Department for providing computational infrastructure and support, as well as Wolfgang Huber for discussions on computing image based scores and statistical tests. We thank the patients who participated in this study. We also thank Christian Drosten at the Charité, Berlin and the European Virus Archive (EVAg) for the provision of the SARS-CoV-2 strain BavPat1. Individual images used in the Fig. 1A courtesy of medical illustrations database—https://smart.servier.com/. This work was in part supported by the Deutsche Forschungsgemeinschaft (DFG, German Research Foundation) – Projektnummer 240245660 – SFB 1129 project 5 (HGK), project 6 (BM), project 11 (RB), project 14 (SB), project 18 (MG) and project Z4 (FH) and by the Deutsches Zentrum fuer Infektionsforschung (VL: project TTU 04.705; RB: project TTU 05.705). SB is supported by the Heisenberg program (project number 415089553) and MLS is supported by the DFG (project number 41607209). The funders had no role in study design, data collection, interpretation, or the decision to submit the work for publication.

## Conflict of Interest

The authors declare they have no conflicts of interest.

## Author Contributions

Model system development: M. Cortese, B. Lucic, B. Cerikan, C.J. Neufeldt, M. Lusic, S. Boulant, M. Stanifer, R. Bartenschlager

Microscopy development: S. Olberg, V. Laketa

Image analysis development: C. Pape, R. Remme, A. Wolny, S. Wolf, L. Cerrone, S. Klaus, M. Ganter, F.A.Hamprecht, A. Kreshuk, C. Tischer, V. Laketa

ELISA assay: S. Wolf, P. Schnitzler

Sera selection and processing: S. Ullrich, M. Anders-Össwein, B. Müller, P. Schnitzler, U. Merle

Data interpretation: C. Pape, R. Remme, A. Wolny, S. Wolf, F. A. Hamprecht, A. Kreshuk, C. Tischer, H-G. Kräusslich, B. Müller, V. Laketa

Study design: F. A. Hamprecht, H-G. Kräusslich, B. Müller, V. Laketa

Manuscript writing, figures, tables: C. Pape, R. Remme, A. Wolny, S. Wolf, L. Cerrone, F. A. Hamprecht, A. Kreshuk, C. Tischer, B. Müller, V. Laketa

All authors have read and approved the final version of the manuscript.

## Notes

### Competing Interest Statement

The authors have declared no competing interest.

### Summary of Updates

abstract slightly modified, references updated, legend of the table 1 extended, text edits

