## Supplementary Figures for "Microscopy-based assay for semi-quantitative detection of SARS-CoV-2 specific antibodies in human sera"

Figure S1

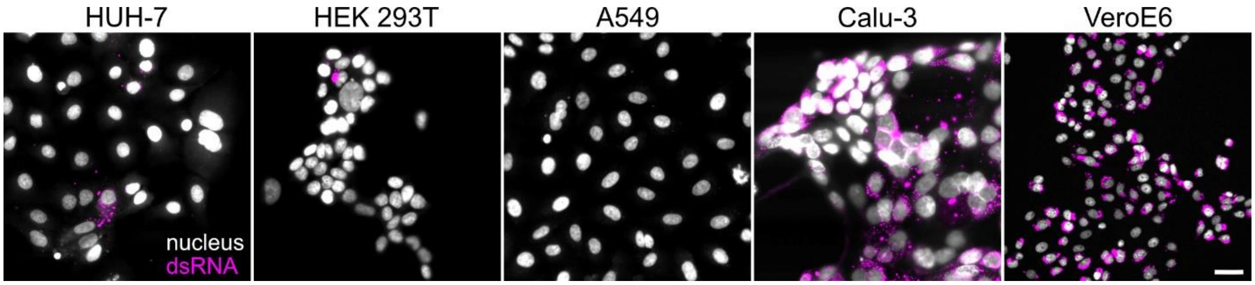


**Figure S1. Different cell lines tested for susceptibility to infection with SARS-CoV-2**. Infection, immunostaining with anti dsRNA antibody and imaging was performed as described in materials and methods. Panels are showing 2-channel overlays of signals for nuclei (Hoechst staining, grey) and dsRNA (magenta) signals for different cell lines. Scale bar = 20 $\mu$m.

Figure S2

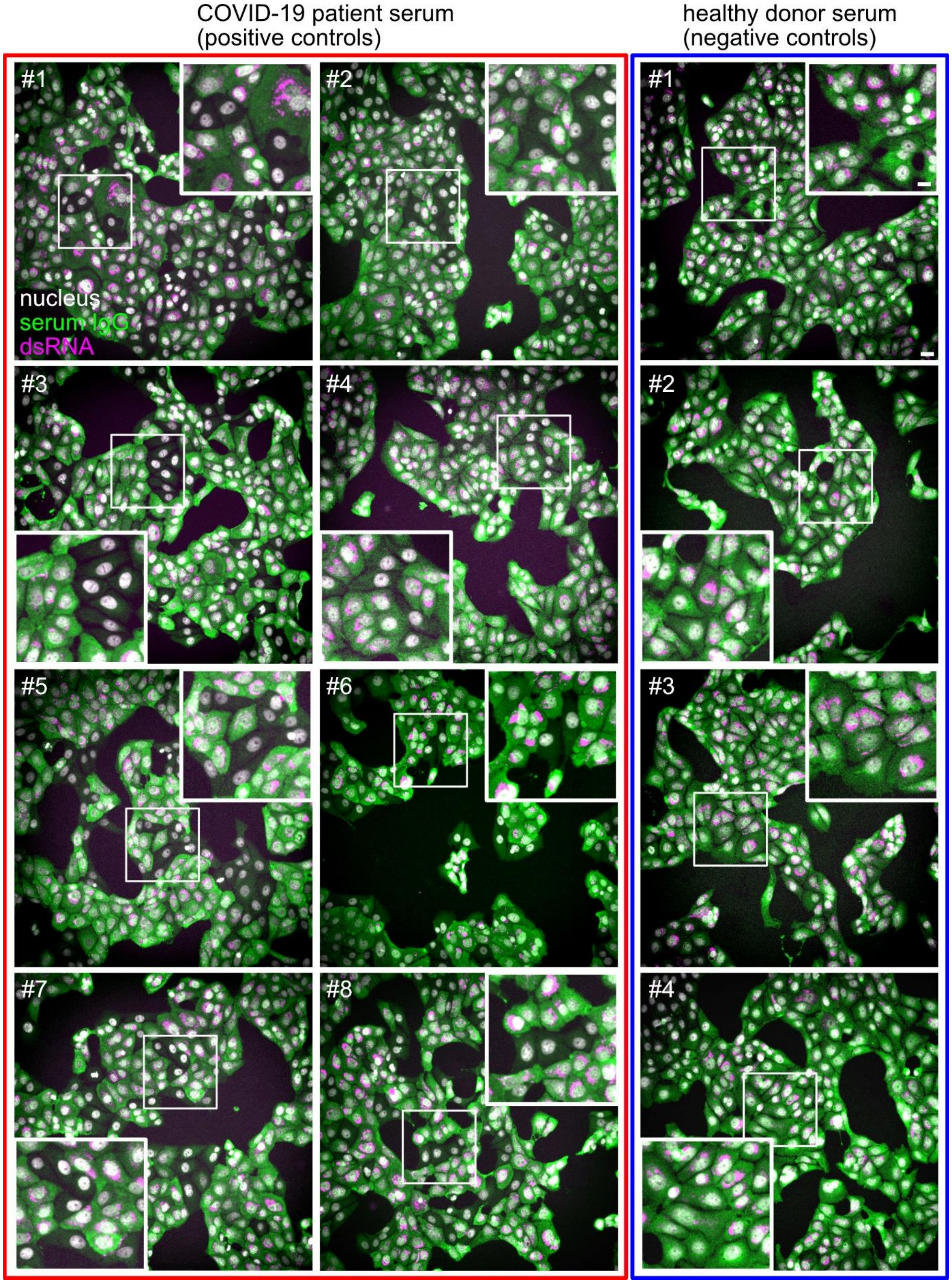
COVID-19 patient serum
(positive controls)
healthy donor serum
(negative controls)
#1
#2
#3
#4
#5
#6
#7
#8
nucleus
serum IgG
dsRNA
#1
#2
#3
#4

**Figure S2. IgG binding to SARS-CoV-2 infected cell samples immunostained with positive and negative control sera**. Panels are showing overlays (nuclei (grey), serum IgG reactivity (green) and dsRNA immunostaining (magenta)) of representative microscopy images of cells stained with eight (#1 - #8) COVID-19 patient sera (positive controls, shown on the left, red box) and four (#1 - #4) sera from healthy donor (pre -COVID-19) serum (negative controls, shown on the right, blue box). White boxes mark the zoomed areas. Scale bar is 20 μm in overviews and 10 μm in zoomed images.

Figure S3

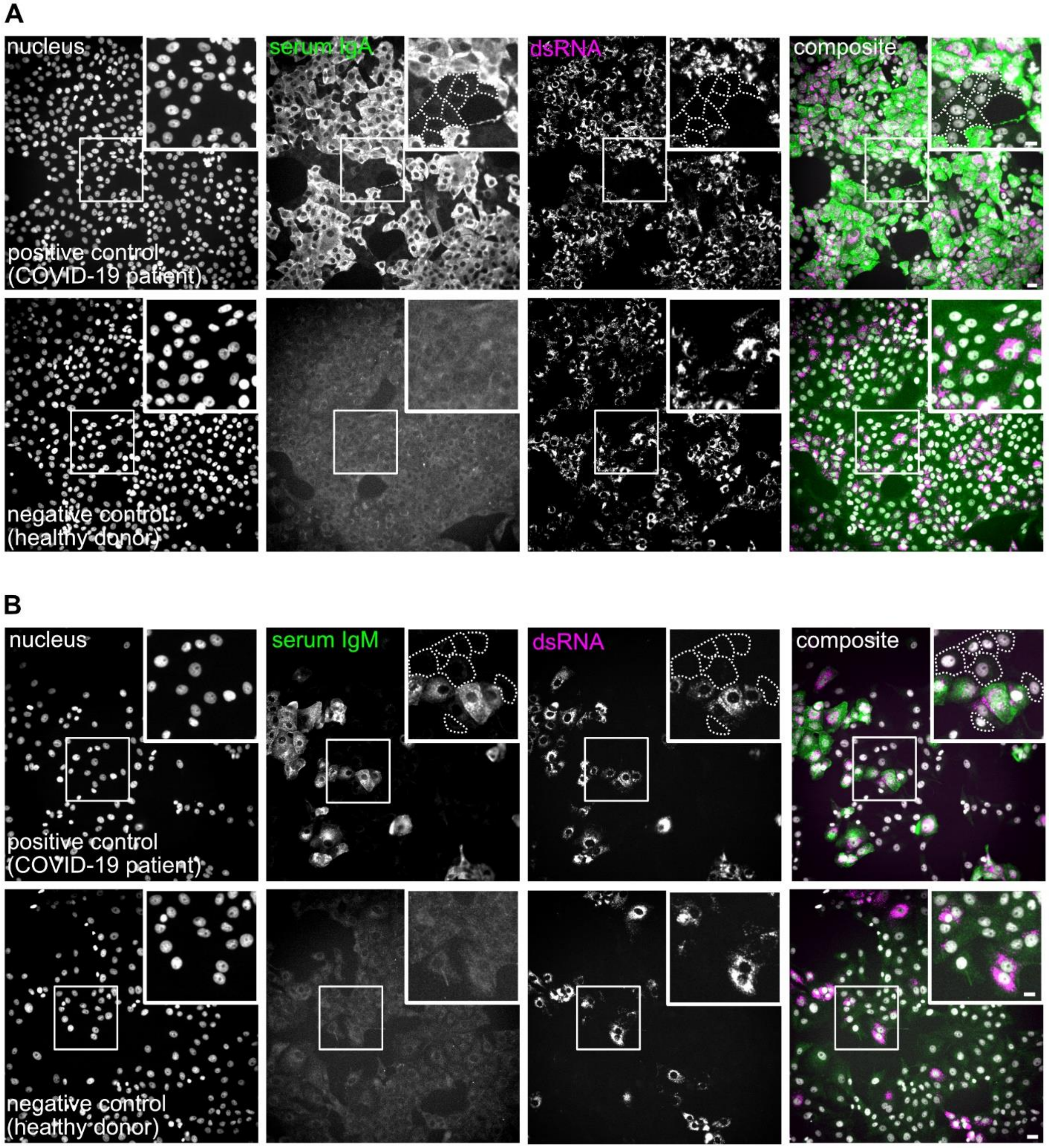


**Figure S3. Assessment of SARS-CoV-2 IgA and IgM specific immunofluorescence staining of infected cell samples stained with positive and negative control sera**. Images show results obtained using the indicated primary antisera and IgA (A) or IgM (B) specific secondary antibodies. Nuclei (grey), IgA or IgM antibody (green), dsRNA (magenta) channels and a composite image are shown. White boxes mark the zoomed area. Dashed lines mark borders of non-infected cells which are not visible at the chosen contrast setting. Note that different brightness and contrast settings were chosen for lower panels in order to visualize cells in the IgA/IgM channel where staining only slightly above the background was detected. Scale bar is 20 μm in overviews and 10 μm in the insets.

Figure S4

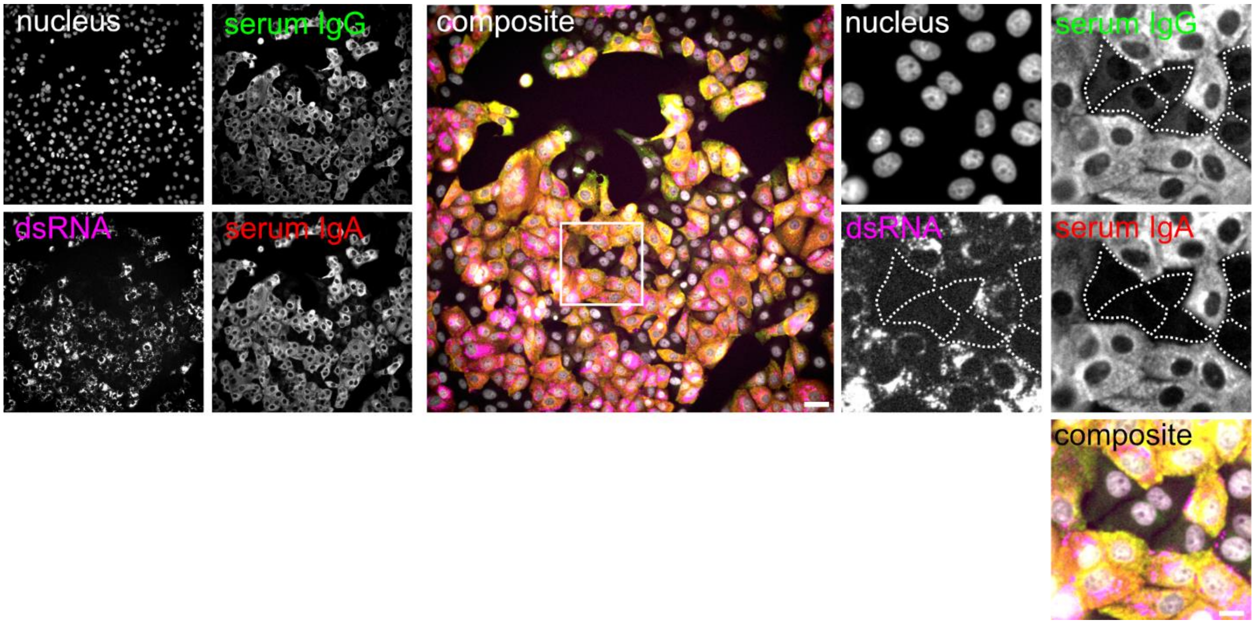


**Figure S4. Simultaneous detection of SARS-CoV-2 IgG and IgA specific staining using COVID-19 patient serum by immunofluorescence**. Nucleus (grey), IgG (green), IgA (red), dsRNA (magenta) cannels and a composite image are shown. White boxes mark the zoomed area. Dashed lines mark borders of non-infected cells which are not visible at the chosen contrast setting. Scale bar is 20 µm in overviews and 10 µm in zoomed images.

Figure S5

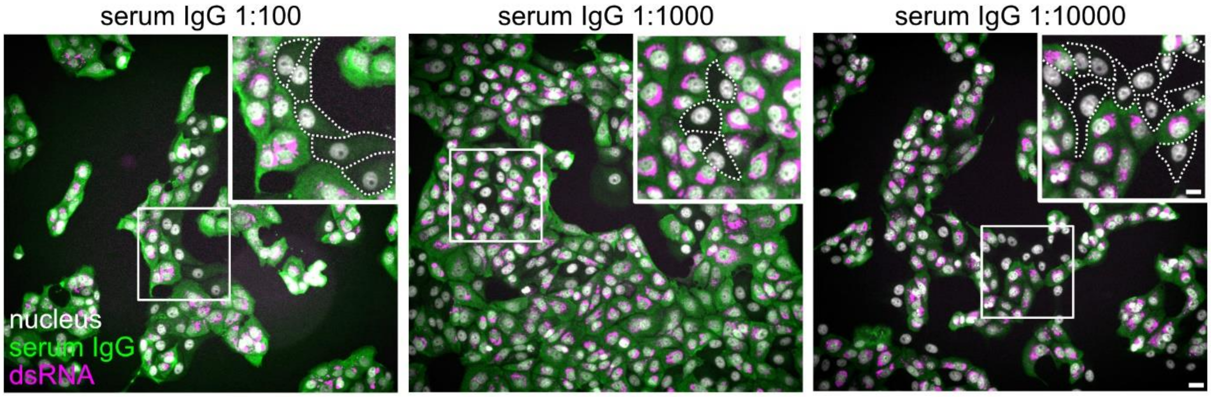


**Figure S5. SARS-CoV-2 IgG immunostaining using serial dilutions of positive control serum by immunofluorescence**. Serum from COVID-19 patient #1 (Supplementary figure 2) was used at the indicated dilution for immunostaining, followed by anti-IgG secondary antibody coupled to AlexaFluor488. Panels display overlay images (nucleus (grey), serum IgG (green) and dsRNA (magenta)) of representative microscopy images. All images were recorded and displayed with identical brightness and contrast settings. White boxes mark the zoomed area. Scale bar is 20 μm in overviews and 10 μm in insets.

Figure S6

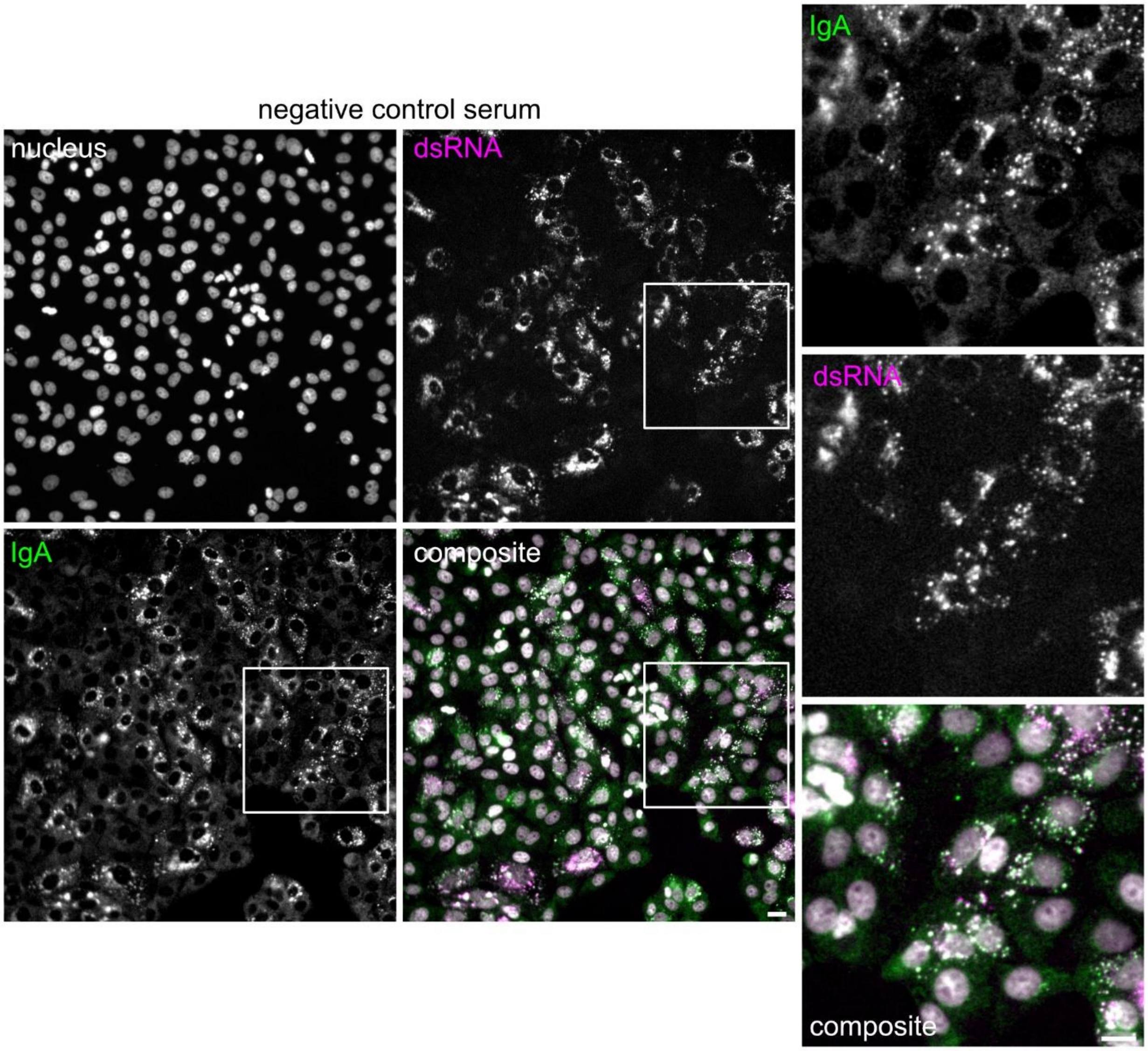


**Figure S6. Nonspecific IgA staining pattern filtered out in a final quality control step**. Images display infected call samples stained with a negative control serum and secondary antibody against IgA. Nucleus (grey), serum IgA (green), dsRNA (magenta) channels and a and composite image are shown. White boxes mark the zoomed area. Note perfectly co-localizing pattern in IgA and dsRNA channels. Scale bar is 20 μm in overviews and 10 μm in zoomed images.

Figure S7

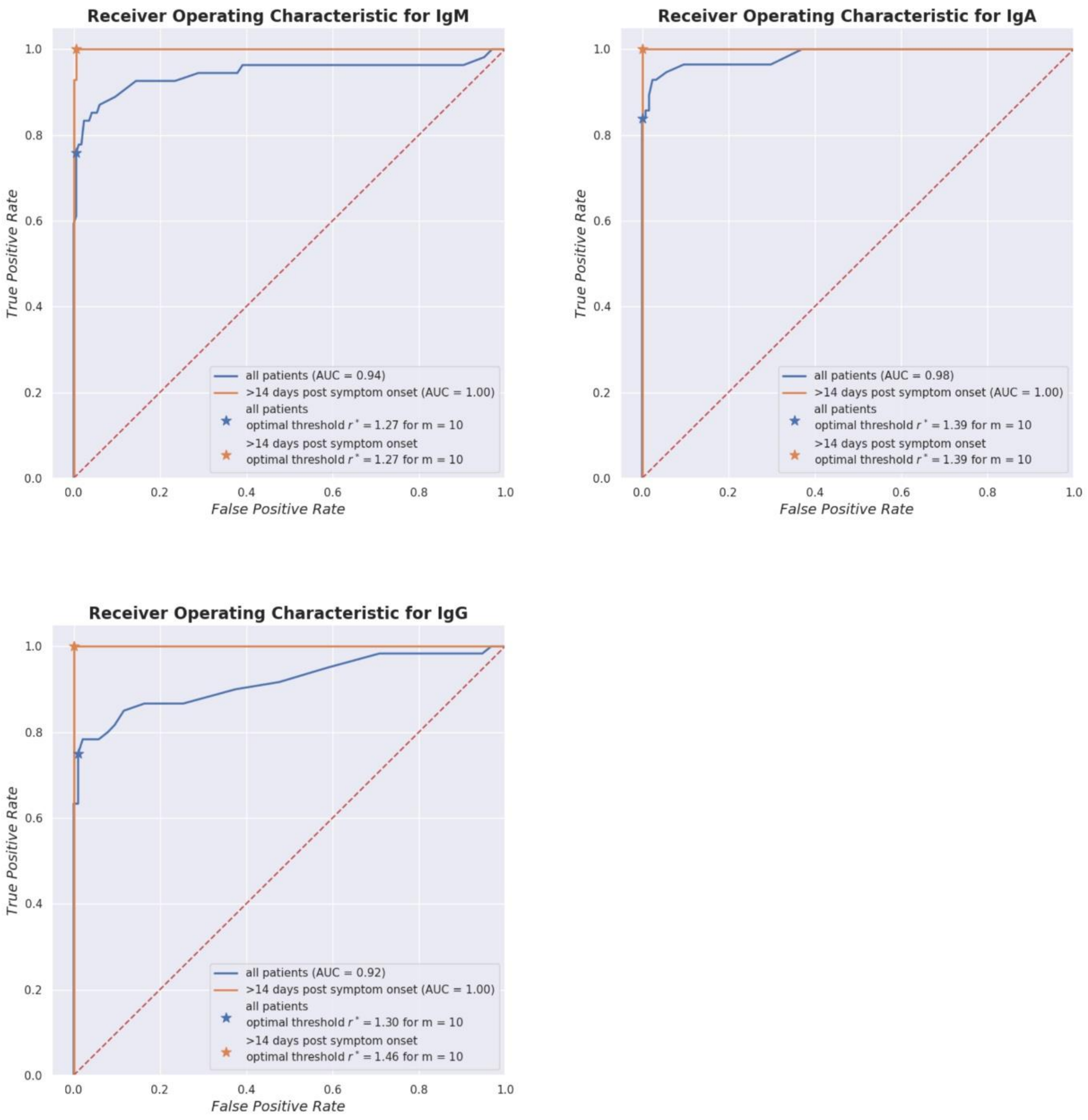


**Figure S7. ROC plot for immunofluorescence (IgM, IgG and IgA) assays.** Solid lines show all possible pairs of (false-positive rate, true-positive rate) or equivalently (1 - specificity, sensitivity) derived from varying the tests decision threshold. Stars show the optimal threshold for our choice of prevalence and costs that corresponds to a slope of m= (false positive costs * (1 - P)) / (false negative costs * P) = 10 (see main manuscript text). Discriminating only between sera from negative patients and sera from patients collected later than d14 post symptom onset improves the accuracy of the test.
